# Parallel Evolution of Linezolid Resistant *Staphylococcus aureus* in Patients with Cystic Fibrosis

**DOI:** 10.1101/2023.05.02.539145

**Authors:** Nicholas J. Pitcher, Andries Feder, Nicholas Bolden, Christian F. Zirbes, Anthony J. Pamatmat, Linda Boyken, Jared J. Hill, Andrew L. Thurman, Valérie C. Reeb, Harry S. Porterfield, Ahmed M. Moustafa, Paul J. Planet, Anthony J. Fischer

**Affiliations:** Stead Family Department of Pediatrics, University of Iowa Carver College of Medicine, Iowa City, IA 52242; Children’s Hospital of Philadelphia, Philadelphia, PA 19104; Perelman School of Medicine, University of Pennsylvania, Philadelphia, PA 19104; Pathology. University of Iowa Carver College of Medicine, Iowa City, IA 52242; Internal Medicine, University of Iowa Carver College of Medicine, Iowa City, IA 52242; State Hygienic Laboratory at the University of Iowa, Coralville, IA 52241; Microbiology Service, Department of Laboratory Medicine, Clinical Center, National Institutes of Health, Bethesda, Maryland, USA; Comparative Genomics, American Museum of Natural History, New York, NY 10024

**Keywords:** *Staphylococcus aureus*, Linezolid, MRSA, ribosomal RNA, hypermutation

## Abstract

**Background:** Linezolid is an antibiotic used to treat serious *Staphylococcus aureus* infections. Resistance to linezolid is considered rare but could emerge with repeated dosing. We recently reported widespread prescription of linezolid for a cohort of patients with cystic fibrosis (CF).

**Objectives:** The goals of this study were to determine the incidence of linezolid resistance in CF and identify molecular mechanisms for linezolid resistance.

**Methods:** We identified patients with *S. aureus* resistant to linezolid (MIC > 4) at the University of Iowa CF Center between 2008 and 2018. We obtained isolates from these patients and retested susceptibility to linezolid using broth microdilution. We used whole genome sequencing to perform phylogenetic analysis of linezolid resistant isolates and examine sequences for mutations or accessory genes that confer linezolid resistance.

**Main Results:** Between 2008 and 2018, 111 patients received linezolid and 4 of these patients cultured linezolid resistant *S. aureus*. We sequenced 11 resistant and 21 susceptible isolates from these 4 subjects. Phylogenetic analysis indicated that linezolid resistance developed in ST5 or ST105 backgrounds. Three individuals had linezolid resistant *S. aureus* with a G2576T mutation in 23S rRNA. One of these subjects additionally had a *mutS^-^ mutL^-^* hypermutating *S. aureus* that produced 5 resistant isolates with multiple ribosomal subunit mutations. In one subject, the genetic basis for linezolid resistance was unclear.

**Conclusions:** Linezolid resistance evolved in 4 of 111 patients in this study. Linezolid resistance occurred by multiple genetic mechanisms. All resistant strains developed in ST5 or ST105 MRSA backgrounds.

**Key Point:** Linezolid resistance arises through multiple genetic mechanisms and could be facilitated by mutator phenotypes. Linezolid resistance was transient, possibly due to growth disadvantage.

## Introduction

*Staphylococcus aureus* is the most prevalent pathogen affecting patients with cystic fibrosis (CF) in the United States (1). *S. aureus* infections are associated with increased airway inflammation in children with CF and precede worsening lung function (2, 3). Infections with *S. aureus* are difficult to eliminate, even with inhaled antibiotics (4). These durable infections persist indefinitely within patients, even after patients acquire *Pseudomonas aeruginosa* infections (5).

Methicillin resistant *Staphylococcus aureus* (MRSA) infections are especially persistent in patients with CF (5). Because MRSA infections are associated with clinical worsening (3, 6), patients with MRSA receive intensified care, including frequent courses of antibiotics (7). Orally bioavailable antibiotics for MRSA include trimethoprim-sulfamethoxazole, tetracyclines, and oxazolidinones. Linezolid is an oxazolidinone antibiotic that is clinically effective in the treatment of severe *Staphylococcus aureus* respiratory infections (8), and it is commonly used in children and adults with CF (9).

Linezolid inhibits ribosomal protein synthesis and is broadly active against Gram-positive bacteria. It binds with high affinity to the ribosomal peptide-transferase center on the 50S subunit. This activity affects tRNA positioning and prohibits protein synthesis (10–12). Although resistance to linezolid is rare (13, 14), clinical linezolid-resistant bacteria have been reported shortly after the drug was introduced (15). Some linezolid resistant bacteria carry mutations in domain V of the 23S rRNA genes, the most common of which is a G to T transversion at position 2576 according to the *Escherichia coli* numbering system (16). *Staphylococcus aureus* encodes multiple copies of the 23S rRNA gene. Thus, there is a gene dosage effect whereby the linezolid MIC increases with the number of mutant copies (17). Resistance to linezolid can also be conferred by acquisition of *cfr*, which encodes a methyltransferase that methylates the 23S rRNA and is often carried on mobile genetic elements. In the absence of *cfr* or 23S rRNA mutations, resistance to macrolide, lincosamide, streptogramin, ketolide, and oxazolidinone (MLSKO) antibiotics has been reported in association with mutations of 50S ribosomal proteins or modifiers such as ribosomal proteins L4 and L22 (18).

Because MRSA is prevalent in patients with CF in the US, repeated prescription of linezolid for chronic unremitting infections creates ideal conditions for evolved linezolid resistance. Additionally, as MRSA is a pathogen of global concern, it is increasingly important to understand how this organism develops resistance to critical antibiotics like linezolid. In this study, we aimed to determine the incidence of linezolid resistance in CF and determine the molecular mechanisms of resistance in this population.

## Materials and Methods

### Ethics statement

The Institutional Review Board (IRB) of the University of Iowa approved study 201905718, a retrospective study of *S. aureus* infections in CF. A waiver of informed consent requirement was granted because the study was minimal risk.

### Subject selection and clinical information

All subjects were patients with CF who received treatment at the University of Iowa CF center between 2008 and 2018. The diagnosis of CF was established by a positive sweat chloride test or genetic testing with two pathogenic mutations in *CFTR*. We excluded any subjects who did not have an electronic prescription or did not have a clinical microbiology report during the observation period. The primary exposure of interest was receiving an electronic prescription or inpatient order for linezolid. Electronic prescriptions became standard of care within this center after February 2009. Linezolid exposure was defined by an electronic order for linezolid by either oral or IV route between February 2009 and April 2018. We recorded the indication for linezolid from each electronic prescription to determine whether it was intended for treatment of *S. aureus* or for other CF pathogens such as non-tuberculous *Mycobacteria*. We compared the prescription dates for linezolid versus other anti-Staphylococcal medications to determine which medication(s) were used to treat infection and in which sequence. Clinical information about these patients, including lung function testing and other outcomes, was recorded to determine the severity of lung disease and rate of complications. We analyzed clinical microbiology reports to determine positivity for relevant CF pathogens, including *S. aureus* and *P. aeruginosa*. The primary outcome of interest was the development of linezolid resistant *S. aureus*.

### S. aureus isolates

The University of Iowa clinical microbiology lab routinely cryopreserved *S. aureus* isolates between 2009 and 2018. We requested any isolates that were cultured from patients who had at least one reported linezolid resistant *S. aureus* on a clinical microbiology report. We confirmed linezolid resistance by broth microdilution antimicrobial susceptibility testing in Mueller-Hinton broth. We selected every isolate from individuals with at least one confirmed linezolid resistant *S. aureus* culture, defined as MIC > 4 µg/mL by broth microdilution, then analyzed isolates by whole genome sequencing (WGS). Of the 111 patients screened, we confirmed that 4 subjects had at least one linezolid resistant *S. aureus*, including 11 linezolid resistant isolates. A total of 32 isolates, 11 resistant and 21 suscepbible, were available for analysis by WGS.

### Whole Genome Sequencing

To determine whether there was transmission of linezolid resistant *S. aureus* between patients and to determine the molecular mechanism for resistance, all isolates were analyzed by WGS as previously described (19). DNA was isolated robotically at the University of Iowa State Hygienic Laboratory.

### Genome Sequencing

We used Illumina DNA Prep and IDT for Illumina DNA/RNA UD indexes to prepare libraries and performed 2x300 bp paired-end sequencing using the Illumina MiSeq Reagent v.3Kit. To produce closed assemblies, we selected 6 isolates for additional long read sequencing using the Rapid Barcoding Kit on a MinION Mk1C device (Oxford Nanopore). We created a hybrid assembly with short and long-read sequences using Unicycler (20). To visualize 23S rRNA alignment, we used MegAlign Pro (DNASTAR) and used *Escherichia coli* strain ATCC 8739 (GenBank accession NC_010468) to indicate position G2576 (21).

### Phylogenetic Analysis

We constructed a maximum likelihood tree for 68 genomes; 32 genomes from our collection and 36 assembled genomes available on GenBank (22), which were the top 5 genomes found using the topgenome (-t) feature of WhatsGNU (23). We processed the genomes from our collection using the Bactopia v1.6.1 (24), and de novo assembly was completed using Shovill v1.1.0 (25). Sequence types were re-analyzed using mlst v2.19.0 (26) which made use of the PubMLST typing schemes (27). We determined the clonal complex for each genome using the WhatsGNU report. ABRicate (28) was used to determine the presence of *mecA* using a database containing genes from TCH1516 (GenBank Assembly GCF_000017095.1).

We annotated genomes using Prokka v1.14.6 (29). A pangenome alignment produced by Roary v3.13.0 (30) was used to infer an initial phylogenetic tree in RAxML v8.2.9 (31). We used a GTR substitution model (32) to account for among-site rate heterogeneity using the Γ distribution and four rate categories (GTRGAMMA model) (33) for 100 individual searches with maximum parsimony random-addition starting trees. Node support was evaluated by running 100 nonparametric bootstrap pseudoreplicates (34). Pairwise SNP distances were calculated using SNP-dist (35).

To optimize visualization, we edited the phylogenetic tree using iTol website (v6.4.2) (36). The data used in this publication were collected through the MENDEL high performance computing (HPC) cluster at the American Museum of Natural History. This HPC cluster was developed with National Science Foundation (NSF) Campus Cyberinfrastructure support through Award #1925590.

### Growth Comparison

To compare the growth of linezolid susceptible versus linezolid resistant isolates, we inoculated isolates in tryptic soy broth (TSB) and incubated with shaking in a 37 °C incubator. We assessed growth using an OD_600_ and recorded the results for the linezolid resistant and susceptible isolates from the same patients. For statistical testing, we used a 2-way ANOVA with pairwise comparisons at each time point.

## Results

### Prescription of Linezolid in Cystic Fibrosis

We surveyed electronic prescriptions and inpatient orders to identify patients receiving linezolid. 360 patients with CF (with or without lung transplant) had encounters between 2009 and 2018, 346 had both medication information and microbiology results. Of these patients, 111 (32%) were treated with linezolid, Supplemental Data 1 and 2. We found 491 oral prescriptions and 343 intravenous prescriptions, consistent with mixed inpatient and outpatient use of linezolid.

### Patient characteristics

We compared patients receiving linezolid to control patients who received other antibiotics. Although it treats other pathogens such as non-tuberculous mycobacteria (37), linezolid treatment was strongly associated with MRSA and other antibiotics targeting MRSA Table 1. MRSA infections are associated with increased age and poorer outcomes in CF (3, 6, 38). We found that patients prescribed linezolid were older and more likely to have died or required lung transplant than controls. These patients also were more likely to have coexisting infections with *P. aeruginosa* and other CF-associated Gram-negative pathogens that are common with advanced lung disease.

**Table 1.** Clinical characteristics of patients prescribed linezolid therapy from 2009 – 2018.

| Characteristic | Linezolid Prescribed | Linezolid Not Prescribed | *P* |
| --- | --- | --- | --- |
| **Number** | 111 | 235 |  |
| **Sex – Female** | 56 (50.5%) | 109 (46.4%) | 0.49 |
| **Genotype** |  |  | 0.08 |
| F508del/F508del | 66 (59.5%) | 112 (47.7%) |  |
| F508del/other | 34 (30.6%) | 101 (43.0%) |  |
| other/other | 11 (9.9%) | 22 (9.4%) |  |
| **CFTR Modulator Therapy** |  |  |  |
| Ivacaftor | 6 (5.4%) | 23 (9.8%) | 0.21 |
| Ivacaftor, Lumacaftor/Ivacaftor, or Tezacaftor/Ivacaftor | 40 (36.0%) | 81 (34.5%) | 0.81 |
| **Anti-Staphylococcal Antibiotics** |  |  |  |
| Trimethoprim-Sulfamethoxazole | 93 (83.8%) | 138 (58.7%) | <0.001 |
| Tetracycline, Doxycycline, or Minocycline | 68 (61.3%) | 52 (22.1%) | <0.001 |
| Vancomycin IV | 96 (86.5%) | 47 (20.0%) | <0.001 |
| Vancomycin Inhaled | 14 (12.6%) | 0 | <0.001 |
| **Birth year, median [IQR]** | 1992<br>[1982-2000] | 1997<br>[1983 – 2009] | 0.002<sup>†</sup> |
| **Outcome\*** |  |  | < 0.001 |
| Death without lung transplant | 10 | 14 |  |
| Death following lung transplant | 14 | 4 |  |
| Alive with lung transplant | 14 | 4 |  |
| Alive without lung transplant, culture after Jan 1 2017 | 62 | 170 |  |
| No known death or transplant, data missing after Jan 1 2017 | 11 | 43 |  |
| **Pathogen Positivity, 2008-2018** |  |  |  |
| MRSA | 94 (84.7%) | 58 (24.7%) | < 0.001 |
| MSSA | 79 (71.2%) | 170 (72.3%) | 0.90 |
| *P. aeruginosa* | 103 (92.8%) | 160 (68.1%) | < 0.001 |
| Mucoid *P. aeruginosa* | 77 (69.4%) | 92 (39.1%) | < 0.001 |
| *Burkholderia spp.* | 13 (11.2%) | 13 (5.5%) | 0.05 |
| *Stenotrophomonas maltophilia* | 58 (52.3%) | 90 (38.3%) | 0.02 |
| *Achromobacter xylosoxidans* | 18 (16.2%) | 16 (6.8%) | 0.01 |

\*Categorical outcomes are as of Dec 31, 2018. *P* values calculated by Fisher's exact test for categorical variables and <sup>†</sup>Wilcoxon rank sum test for birth year.

### Linezolid prescriptions were given to treat *S. aureus* infections and followed other attempts to treat MRSA

The indication for linezolid was written for 197 (40%) of the oral prescriptions. The most common indication for linezolid was treatment of *S. aureus* or MRSA (N = 131), followed by treatment of CF, bronchitis, or pulmonary exacerbation (N = 57) followed by non-tuberculous Mycobacteria (N = 4).

Three other antibiotic classes are commonly used to treat MRSA in CF: vancomycin, trimethoprim-sulfamethoxazole, and tetracyclines (38). In our cohort, 53 patients were treated with all four classes. Within these patients, trimethoprim/sulfamethoxazole was used first, followed by vancomycin, then linezolid and tetracyclines, <u>Supplemental Data 3</u>. These data suggest that linezolid was usually reserved for later attempts at treating MRSA.

### Repeated prescription of linezolid

Repeated linezolid dosing in patients with chronic infection may increase the risk of evolved resistance. Of the 111 subjects receiving linezolid 27 subjects received only one prescription, whereas 62 patients received 5 or more prescriptions, Supplemental Data 4. The maximum number of linezolid prescriptions written for a single patient was 50.

### Resistance of *S. aureus* to linezolid

Linezolid resistant *S. aureus* were reported in 11 patient encounters for 5 patients. We obtained *S. aureus* isolates, including linezolid susceptible and resistant, from these patients and performed antibiotic susceptibility testing by broth microdilution. We confirmed linezolid resistance (MIC ≥ 8 mg/L) in isolates from 4 of the 5 subjects. These four patients received between 8-20 orders of linezolid, Figure 1. We analyzed 11 linezolid resistant and 21 susceptible isolates by WGS, Table 2.

**Figure 1.**
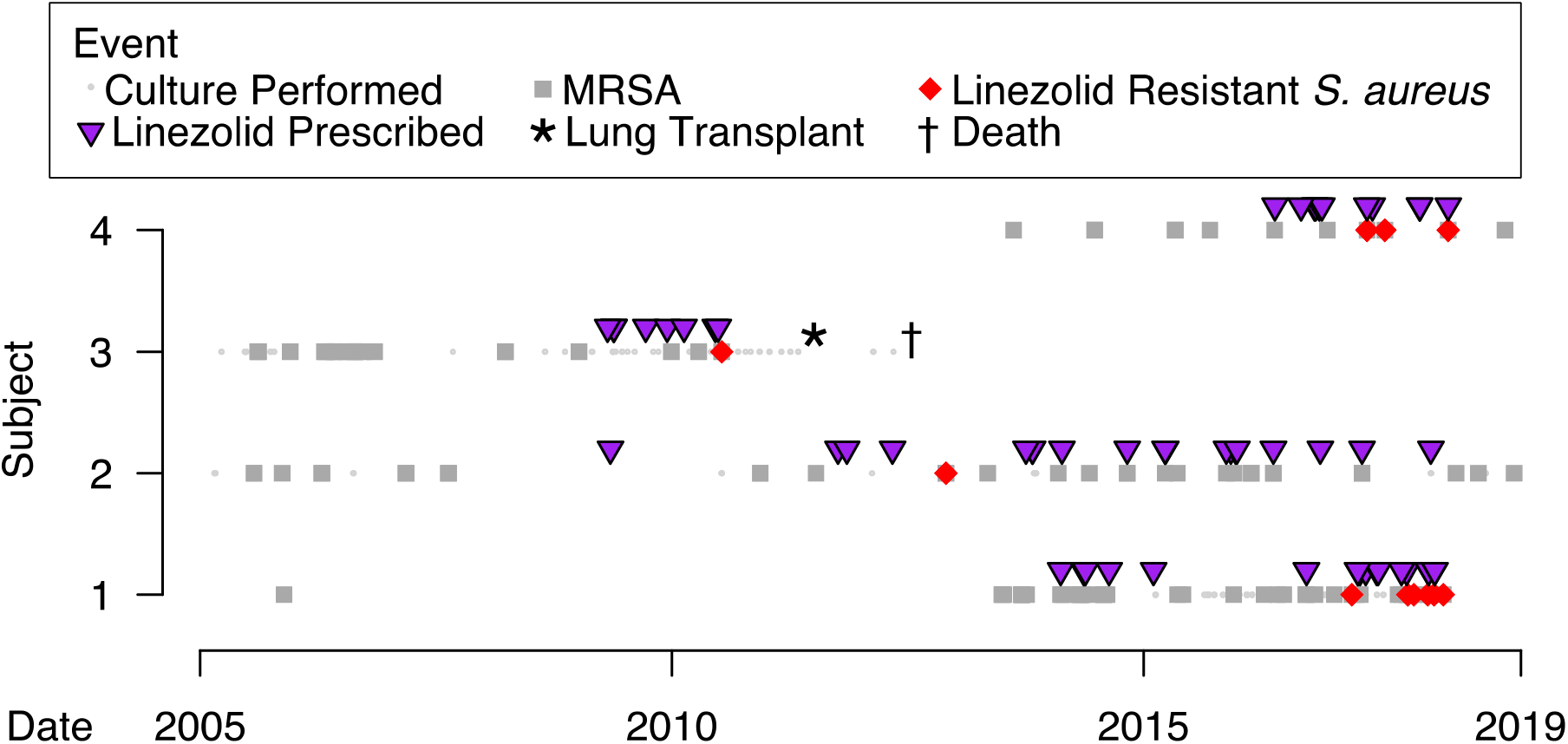
Timelines of *S. aureus* positivity and linezolid prescription in 4 patients with cystic fibrosis who had confirmed linezolid resistant MRSA. Small dots indicate a culture was performed. Dark gray squares indicate a culture positive for MRSA. Red diamonds indicate linezolid resistant *S. aureus*. Purple triangles above the timeline indicate prescriptions for linezolid.

**Table 2.** Isolates analyzed by WGS. Shading indicates linezolid resistance.

| Subject # | Isolate | Collection Year-month | Linezolid MIC | Oxacillin MIC | 23S rRNA copy number* | MLST | Mechanism of Resistance |
| --- | --- | --- | --- | --- | --- | --- | --- |
| Subject #1 | AF4001 | 2015-06 | 4 | 16 | 6 | 5 |  |
|  | HP20814.006 | 2017-01 | 4 | >16 | 6 | 5 |  |
|  | AF2324 | 2018-01 | >8 | >16 |  | 5 | G2576T |
|  | HP20814.043 | 2017-03 | 8 | >16 | 6 | ND | Multiple |
|  | HP20814.051 | 2017-04 | 4 | >16 |  | ND |  |
|  | AF4003 | 2017-10 | 8 | >16 |  | ND | Multiple |
|  | HP20814.091 | 2017-11 | 8 | >16 |  | ND | Multiple |
|  | AF2323 | 2018-01 | 8 | >16 |  | ND | Multiple |
|  | AF2325 | 2018-03 | 8 | >16 |  | ND | Multiple |
| Subject #2 | AF4005 | 2012-11 | 8 | >16 | 5 | 105 | Unknown |
|  | HP20814.058 | 2017-04 | 4 | >16 |  | 105 |  |
|  | AF2246 | 2018-04 | 4 | >16 |  | 105 |  |
|  | AF2247 | 2018-07 | 2 | >16 |  | 105 |  |
|  | AF2248 | 2018-12 | 2 | >16 |  | 105 |  |
| Subject #3 | AF4006 | 2006-04 | 4 | >16 |  | 5 |  |
|  | AF4008 | 2006-06 | 4 | >16 |  | 5 |  |
|  | AF4009 | 2006-06 | 4 | >16 |  | 5 |  |
|  | AF4010 | 2006-07 | 4 | >16 |  | 5 |  |
|  | AF4011 | 2006-08 | 2 | >16 |  | 5 |  |
|  | AF4012 | 2006-09 | 2 | >16 |  | 5 |  |
|  | AF4013 | 2006-11 | 2 | >16 |  | 5 |  |
|  | AF4015 | 2009-01 | 4 | >16 |  | 5 |  |
|  | AF4016 | 2009-12 | 4 | >16 |  | 5 |  |
|  | AF4017 | 2010-04 | 4 | >16 |  | 5 |  |
|  | AF4004 | 2010-07 | 8 | 16 | 5 | 5 | G2576T |
|  | AF4007 | 2006-05 | 4 | >16 |  | 105 |  |
|  | AF4014 | 2008-03 | 4 | >16 |  | 582 |  |
| Subject #4 | AF1552 | 2015-09 | 4 | 0.25 |  | 5 |  |
|  | HP20814.062 | 2017-05 | 8 | >16 | 6 | 5 | G2576T |
|  | AF4002 | 2017-07 | 8 | >16 |  | 5 | G2576T |
|  | AF2001 | 2018-03 | >8 | >16 |  | 5 | G2576T |
|  | AF2002 | 2018-11 | 4 | 4 |  | 5 |  |

\*Number of rRNA copies is based on hybrid assembly using long read (Oxford Nanopore Minion) and short read (Illumina MiSeq) sequences using Unicycler software.

### Whole Genome Sequencing Analysis

All 11 linezolid resistant isolates evolved from either ST5 or ST105 MRSA backgrounds. These clonal complex 5 lineages are usually SCC*mec* II positive and considered hospital-associated MRSA. Collectively, these sequence types are the most prevalent MRSA lineages in our center and are associated with lower lung function in younger patients compared to clonal complex 8 MRSA lineages (19).

A previous report suggested that these hospital-acquired MRSA lineages have greater potential to evolve resistance to protein synthesis inhibitor drugs due to lower rRNA copy number (39). To determine rRNA copies we used hybrid assembly with short and long-read WGS. Two subjects had MRSA with 6 rRNA operons; the other two had MRSA with 5 rRNA operons, Table 3. Thus, linezolid resistance was not limited to strains with lower rRNA copy number.

**Table 3.** A combination of ribosomal variants in linezolid resistant *S. aureus* arising in a mutator background

|  |  | DNA Repair | 50S Ribosome Proteins/Modifiers |  |  |  |  |  |  | 30S Ribosome Proteins/Modifiers |  |  |
| --- | --- | --- | --- | --- | --- | --- | --- | --- | --- | --- | --- | --- |
|  |  | *mutS*, *mutL* | 23S<br>rRNA | *cfr* | *rplC* | *rlmI* | *rluD* | *rlmN* | *rplD* | *rpsC* | *rpsJ* | *rpsD* |
| Isolate | Linezolid<br>MIC | SA_RS06425<br>SA_RS06430 |  |  | SA_RS<br>11765 | SA_RS<br>05285 | SA_RS<br>05915 | SA_RS<br>06020 | SA_RS<br>11760 | SA_RS<br>11735 | SA_<br>RS11770 | SA_RS<br>08675 |
| HP20814.051 | 4 | wt | wt | absent | wt | wt | wt | wt | wt | wt | wt | wt |
| AF4001 | 4 | wt | wt | absent | p.S145del | wt | wt | wt | wt | wt | wt | wt |
| HP20814.006 | 4 | wt | wt | absent | wt | wt | wt | wt | wt | wt | p.K57M | wt |
| AF2325 | 8 | deleted | wt | absent | p.S145del | p.V218G | wt | wt | wt | wt | p.V55A,<br>p.K57N,<br>p.Y58C | wt |
| HP20814.091 | 8 | deleted | wt | absent | p.S145del | p.V218G | p.L158F | wt | wt | wt | p.K57N,<br>p.Y58C | p.L193S |
| HP20814.043 | 8 | deleted | wt | absent | p.S145del | p.V218G | p.L158F | wt | wt | wt | p.K57N,<br>p.Y58C | p.L193S |
| AF2323 | 8 | deleted | wt | absent | p.S145del | p.V218G | p.L158F | p.Q52fs | wt | wt | p.K57N,<br>p.Y58C | p.L193S |
| AF4003 | 8 | deleted | wt | absent | p.S145del | p.V218G | p.L158F | p.Q52fs | wt | wt | p.K57N,<br>p.Y58C | p.L193S |
| AF2324 | >8 | wt | G2576T | absent | wt | wt | wt | wt | wt | wt | p.K57R | wt |

## Assignment of Isolates into Strains by Clade Breaker

Determining whether linezolid resistant MRSA spreads between patients requires a working definition of a strain. For acute outbreaks related to the spread of MRSA isolates taken from different patients may be highly similar (SNP distance < 20) (40). In chronic infections like CF there can be greater diversity within a single patient. Therefore, the first step in analyzing these isolates is to assign them to strains. Some patients with CF are infected by multiple strains of *S. aureus,* and occasionally unrelated patients share the same strain (37). To determine whether there was sharing of linezolid resistant strains between patients, we performed phylogenetic analysis to assign closely related isolates to strains. Because some CF-associated MRSA strains exhibit hypermutation (19) or have increased SNP distance as they develop linezolid resistance (41, 42), we did not use strict SNP cutoffs to divide isolates into strains. Instead, we used a clade-breaker approach (43). For each isolate, we downloaded the five most closely related *S. aureus* genomes from GenBank. If a given isolate from the patient was more closely related to an independent strain from GenBank than it was to another isolate taken from the same patient, we assumed the patient’s infection was polyclonal, as the patient’s two isolates were separated by a “clade-breaker.” Results of the phylogenetic analysis are shown in Figure 2 and <u>Supplemental Data 5</u>. The clade breaker method helped classify several isolates that would have had ambiguous strain assignments using a strict SNP distance relationship alone; several pairwise SNP distances within strains were greater than 60, Supplemental Data 6, 7, 8, 9. Some of the pairwise SNP distances within strains approached the pairwise distances measured between distinct strains isolated from different subjects, which were generally > 200, Supplemental Data 10.

**Figure 2.**
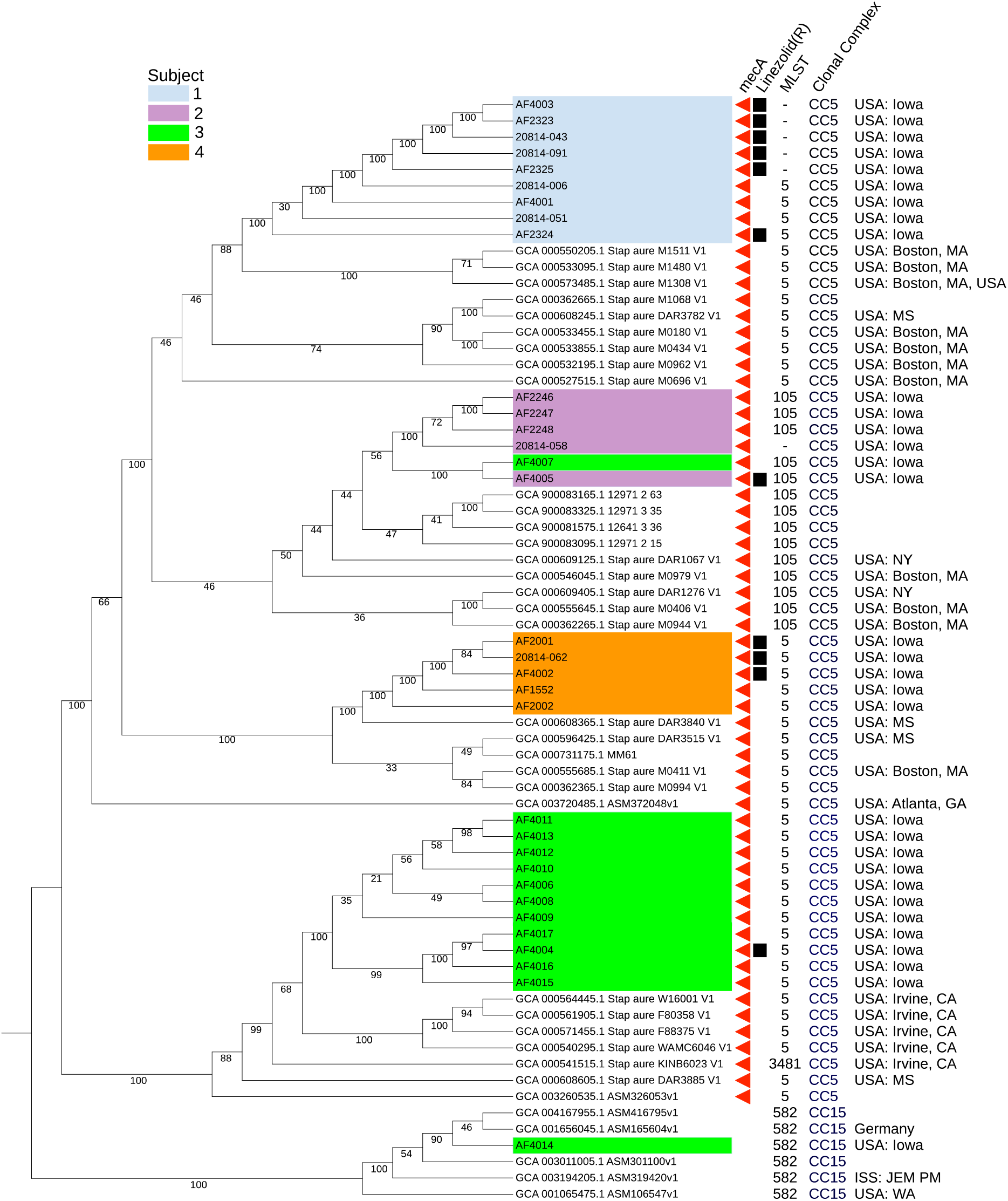
Phylogenetic analysis of Linezolid resistant *S. aureus*. Linezolid resistant isolates (MIC > 4) are indicated by a black square, MRSA are indicated by a red triangle. Isolates analyzed this study (N = 32) are color-coded by study subject. Non-colored branches (N = 36) represent genomes identified from GenBank that were among the top 5 most closely related genomes to those in our study. The tree is a maximum likelihood phylogenetic tree, with numbers underneath each branch of the tree indicating the confidence level of the node based on 100 non-parametric bootstrap pseudoreplicates. Separation of isolates that are shaded with the same color indicates polyclonality of strains cultured from the patient. For example, subject 3 (green) possesses at least three distinct strains. Strain sharing is evident between subject 3 and subject 2 (lavender); the linezolid resistant isolate AF4005 from subject 2 is most closely related to AF4007 from subject 3. All linezolid resistant isolates in this study emerged from ST5 or ST105 MRSA lineages. Within subject 1 (sky blue), phylogenetic analysis suggests linezolid resistance evolved twice, as AF2324 appears separate from the subject’s five other linezolid resistant isolates.

### Linezolid resistance associated with 23S rRNA mutations evolved independently in multiple strains

We previously reported that closely related MRSA were shared between patients (19). Sharing of strains would suggest transmission of resistant organisms between patients. If this were occurring with linezolid resistance, it would threaten linezolid effectiveness across the CF population.

Linezolid resistant *S. aureus* isolates, labeled in Figure 2 with black squares, appeared in 5 separate clusters. Within Subject 1, linezolid resistance evolved twice from the patient’s ST5 ancestor. One of the linezolid resistant isolates (AF2324) diverged from the earliest common ancestor whereas a second series of five linezolid resistant isolates diverged later. Long read sequencing of AF2324 revealed 6 copies of the 23S rRNA operon and confirmed 2 copies of a G2576T mutation known to cause linezolid resistance, Supplemental Data 11. The remaining 5 isolates from this subject had normal 23S rRNA, implying an independent resistance mechanism.

### Linezolid resistance on hypermutator strain background

We examined the remaining linezolid isolates from Subject 1 further. Although they were descended from ST5, these isolates were not assigned a sequence type owing to disruption of the MLST gene *glpF*. Furthermore, short read sequences did not detect the neighboring genes *mutS* and *mutL*, which play important roles in DNA mismatch repair. Drawing the phylogenetic tree with branch lengths proportional to nucleotide polymorphisms, the isolates lacking *mutS* and *mutL* had longer inferred branch lengths, suggesting a hypermutator phenotype, <u>Supplemental Data 5</u>. To determine how these isolates lost *mutS* and *mutL*, we used long-read sequencing to compare isolates with *mutS* and *mutL* (AF4001 and HP20814-006) to an isolate lacking these genes (HP20814-043).

Long read sequencing identified a transposable element with amplified copy numbers in this strain (IS1181). IS1181 was originally absent near *mutS* and *mutL* as shown for AF4001 (Figure 3), but subsequently inserted twice flanking these genes. As shown in HP20814-006, IS1181 inserted within *glpF* and upstream of *mutS*. A subsequent recombination event between these transposable elements resulted in complete excision of *mutS*, *mutL*, *glpP*, and the 5’ portion of *glpF*.

Following the loss of *mutS* and *mutL*, all subsequent isolates were linezolid resistant despite a normal 23S rRNA sequence. Instead, the strain accumulated a series of mutations in genes that encode ribosomal accessory proteins or modifiers, Table 3. These mutations included a previously reported variant p.S145del in *rplC*, as well as new variants in 50S ribosomal genes *rlmI*, *rluD*, and *rlmN*. Additionally, the strain accumulated mutations affecting the 30S ribosomal subunit, which could facilitate resistance to other protein synthesis inhibitors like tetracyclines.

**Figure 3.**
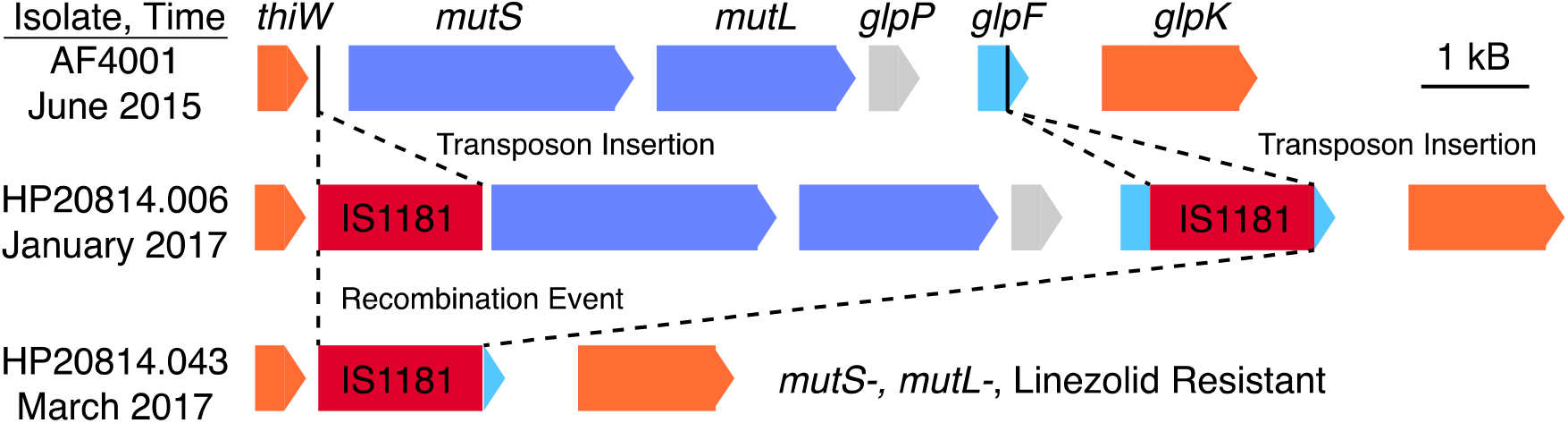
Genetic deletion of DNA mismatch repair genes *mutS* and *mutL* in a strain from subject 1 that developed linezolid resistance. Isolates were taken at different timepoints. *S. aureus* chromosome assemblies from short and long-read sequences are presented for genes near *mutS* and *mutL*. In the second isolate, HP20814.006, transposon IS1181 was inserted twice within the locus, including 5’ of *mutS* and within the coding sequence of *glpF*. A subsequent recombination event deleted *mutS*, *mutL*, *glpP*, and the 5’ coding sequence of *glpF*. In addition to HP20814.043, four other linezolid-resistant isolates lacked *mutS* and *mutL* in this patient.

## Sharing of a Strain Capable of Linezolid Resistance

As reported by others, we observed examples of polyclonality of *S. aureus* and shared strains even in this small patient sample (37). Subject 3 had a polyclonal infection including three distinct *S. aureus* strains, Figure 2. One isolate was ST105 (AF4007), belonging to the same strain infecting Subject 2 (AF4005). AF4005 was linezolid resistant, suggesting that direct or indirect transmission of a strain capable of evolving linezolid resistance occurred between two different patients. These isolates were cultured 6 years apart and it is unclear how this strain was shared. AF4005 had a normal 23S rRNA sequence, and the molecular basis for linezolid resistance remains unclear.

The strain that would ultimately evolve a linezolid resistant isolate in Subject 3 (AF4004) was independent of the strain shared with Subject 2. AF4004 was ST5 and had G2576T mutations in 23S rRNA. Thus, although there was strain sharing between Subject 2 and Subject 3, the evolution of linezolid resistance was phylogenetically and mechanistically distinct in both patients.

Subject 4 had three linezolid resistant isolates, all of which had G2576T mutations. An example from this strain is given in Supplemental Data 12. These isolates belonged to the same strain and were independent of strains obtained from other subjects. Together, resistance to linezolid developed independently in each of the four subjects. G2576T mutations explained linezolid resistance that appeared in three subjects. One subject had linezolid resistance that evolved at least twice during the observation period.

### Growth disadvantage of linezolid-resistant *S. aureus* isolates

Although linezolid resistance should be an advantageous phenotype for *S. aureus*, we were surprised to see that some linezolid resistant strains appeared only transiently despite continued antibiotic pressure. The lack of persistent linezolid resistant *S. aureus* suggests it is at a growth disadvantage compared to linezolid susceptible *S. aureus* (44). Therefore, we compared the growth of linezolid-resistant *S. aureus* to linezolid susceptible *S. aureus* isolated from the same patients, Figure 4. Linezolid susceptible isolates grew significantly better at 4 hours (p < 0.01) and at 6 hours (p < 0.0001) than linezolid resistant isolates, confirming that linezolid resistance comes with a fitness cost, perhaps accounting for low persistence in the CF airway.

**Figure 4.**
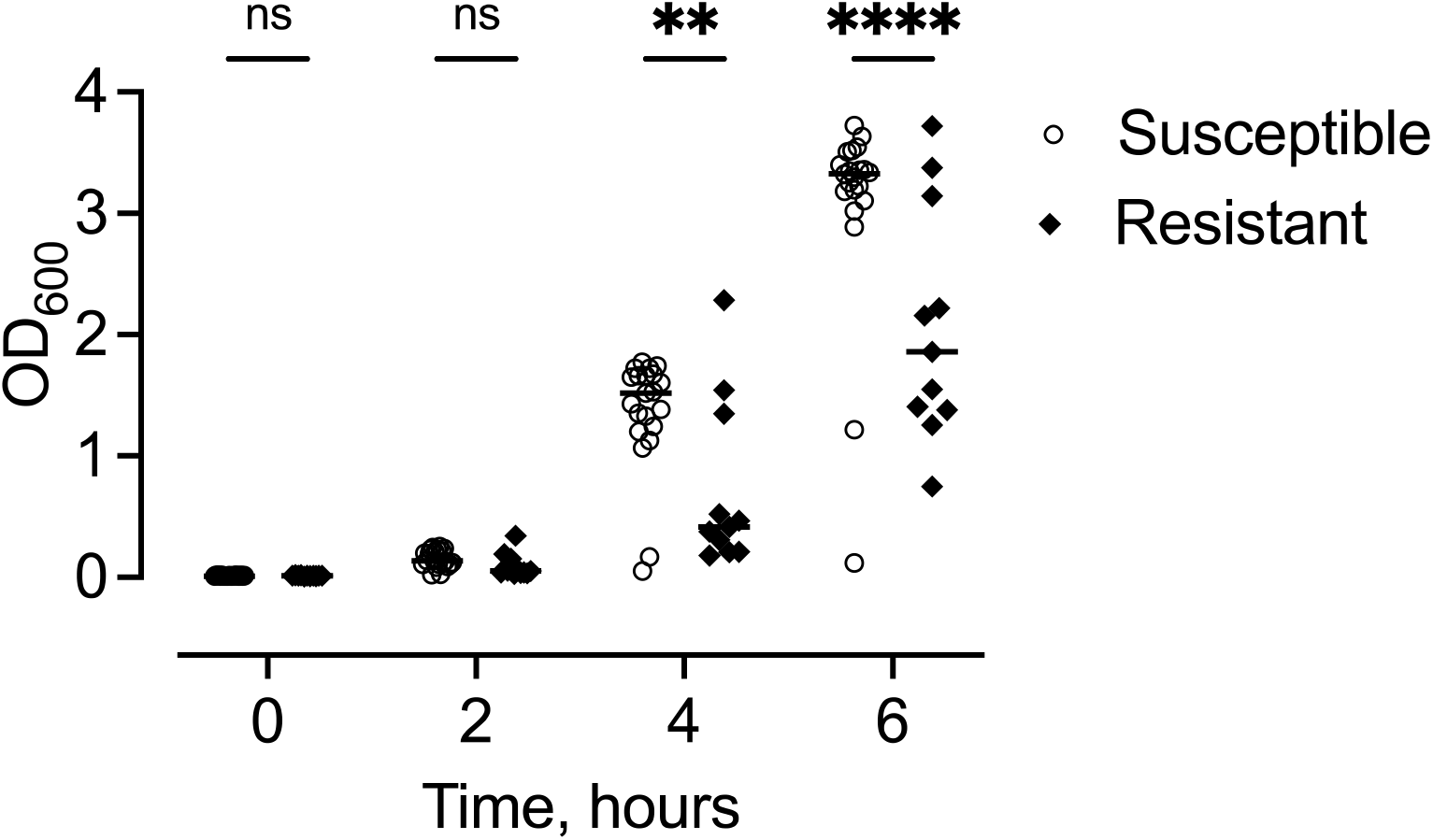
Growth of linezolid susceptible and linezolid resistant *S. aureus* from patients with CF in tryptic soy broth. The growth of linezolid resistant *S. aureus* is slower compared to linezolid susceptible isolates. 2-way ANOVA with pairwise comparisons at each time point. Each dot represents a different isolate and lines indicate the median. N=21 susceptible and 11 resistant isolates ** p < 0.01, **** p < 0.0001, ns = not significant.

## Discussion

We show that of the 111 patients treated with linezolid at the University of Iowa CF center over nearly a decade, 4 developed confirmed linezolid resistant *S. aureus*, an incidence of approximately 4 per 1000 person-years. Subjects who received linezolid often received repeated antibiotic courses because of persistent MRSA. In general, these patients received linezolid after first taking other anti-staphylococcal treatments. These patients had lower lung function than patients who did not receive linezolid. When an indication was provided, linezolid was prescribed more often for treating persistent *S. aureus* infections compared to other CF pathogens like non-tuberculous mycobacteria.

The strains of linezolid resistant *S. aureus* that we examined evolved independently in each patient without acquisition of *cfr*. The most common genetic explanation for linezolid resistance was a well-known G2576T mutation in the 23S rRNA. Notably, one subject evolved linezolid resistance twice by two independent mechanisms. In one strain from this subject, the loss of DNA repair genes *mutS* and *mutL* facilitated a series of mutations in the 50S ribosomal subunit that may have exerted a combinatorial effect on linezolid resistance.

### Comparison to previous studies

The G2576T transversion in 23S rRNA is a frequently reported mechanism of linezolid resistance (15, 42, 45–48). Three patients in this study had *S. aureus* with this mutation, which evolved independently in each case. Mutations in ribosomal L3 protein that contribute to linezolid resistance are rare. Five isolates in a mutator lineage contained a Δser145 variant previously reported in association with linezolid resistance (49). However, expression of this mutation in *E. coli* was not sufficient for linezolid resistance (49). Two of the isolates in this study contained frameshift mutations in the *rlmN* methyltransferase. While this mutation alone is typically not sufficient to confer linezolid resistance, LaMarre and colleagues showed that inactivation of RlmN resulted in increased linezolid MIC (50). It is possible that these mutations act synergistically with other mutations that alter the 50S ribosome to further adapt to linezolid.

Hypermutating bacterial strains can develop resistance to multiple antibiotics (51–53). Ba and others observed that Linezolid resistance developed in *Enterococcus faecalis* mutators, but resistance did not develop in laboratory strains of *S. aureus* with targeted disruption of DNA repair genes (54). It is possible that linezolid resistance did not occur under these settings because evolving resistance may require time to develop a combination of several point mutations. By contrast, antibiotic resistance increased following disruption of *mutL* or *mutS* by frameshift mutations in *P. aeruginosa* (19). Our study shows that linezolid resistant *S. aureus* can evolve in a strain with genetic features of hypermutation, including loss of DNA genes and increased number of SNPs. The mechanism of hypermutation was related to complete loss of *mutL* or *mutS* from a recombination event, which facilitated resistance to linezolid by a series of mutations affecting the 50S ribosome subunit. Insertional inactivation of DNA repair genes has been suggested to have stronger predisposition to antibiotic resistance versus potentially reversible point mutations (55).

### Advantages of our approach

Because we had nearly complete electronic prescription information in stable patient cohort, we had a quantitative measure of prolonged exposure to linezolid and other anti-staphylococcal antibiotics in a population at risk of linezolid resistance. The combined use of short-read and long-read WGS allowed us to determine the number of rRNA operons and to visualize structural rearrangements in the *S. aureus* chromosome. With these resources, we were able to capture changes in the genome outside of the predicted antibiotic targets. Because some strains have hypermutation features, the use of a clade-breaker phylogenetic approach allowed flexibility in determining strain assignments despite occasional large differences in SNPs between isolates collected from the same subject. Longitudinal data collecting, including linezolid susceptible and resistant isolates, allowed us to infer the order of mutations within individual patients and to assess how linezolid resistance evolved.

### Study Limitations

Our study was single-centered, so our findings may not generalize to centers that prescribe less linezolid. Our approach could also underestimate the incidence of linezolid resistance. We studied single isolates per study encounter; because we found that linezolid-resistant *S. aureus* had a growth disadvantage, it is possible that we have underestimated the incidence of linezolid resistance due to under-sampling. We also recently found that *S. aureus* is underdiagnosed in this population due to culture conditions (56). Because the patients studied were treated with a wide range of antibiotics, it is possible that some bacterial mutations are adaptive for antibiotics other than linezolid but confer cross-resistance. Finally, novel mutations that we reported are associated with linezolid resistance, but we have not provided causal evidence of linezolid resistance by analyzing isogenic strains with and without each mutation individually.

## Conclusions

Linezolid is used frequently for patients with CF who have MRSA infections and worsening lung disease. Despite the repeated use of linezolid for some MRSA infections, relatively few subjects developed linezolid resistant MRSA within this center. Persistence of linezolid resistant *S. aureus* within these subjects was uncommon, possibly due to growth defects of linezolid resistant strains. 23S rRNA mutations were the most common cause of linezolid resistant MRSA. Hypermutating strains of MRSA could develop linezolid resistance through a combination of mutations affecting the 50S ribosome. As more molecular diagnostic methods are used in CF, these observations may aid in earlier detection of antimicrobial resistance in patients at risk of developing antibiotic resistance.

### Data Availability

Genome sequencing data has been deposited in GenBank. Accession Numbers for each isolate are provided in Supplemental Data 12. Unassembled sequences are available from the study’s corresponding author. Bacterial strains are available to qualified researchers following completion of a materials transfer agreement.

---

**Supplemental Data 1.**
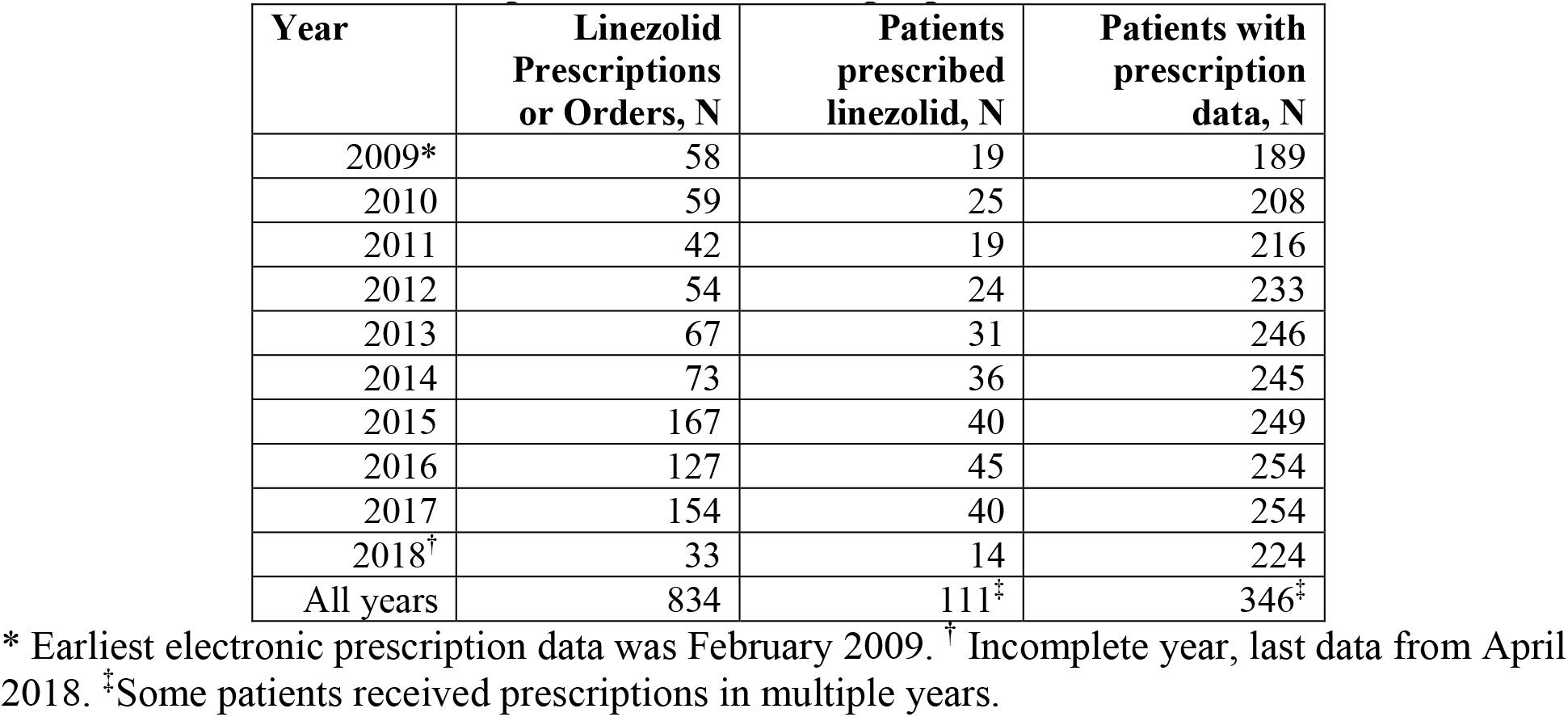
Prescription of Linezolid to people with CF.

**Supplemental Data 2.**
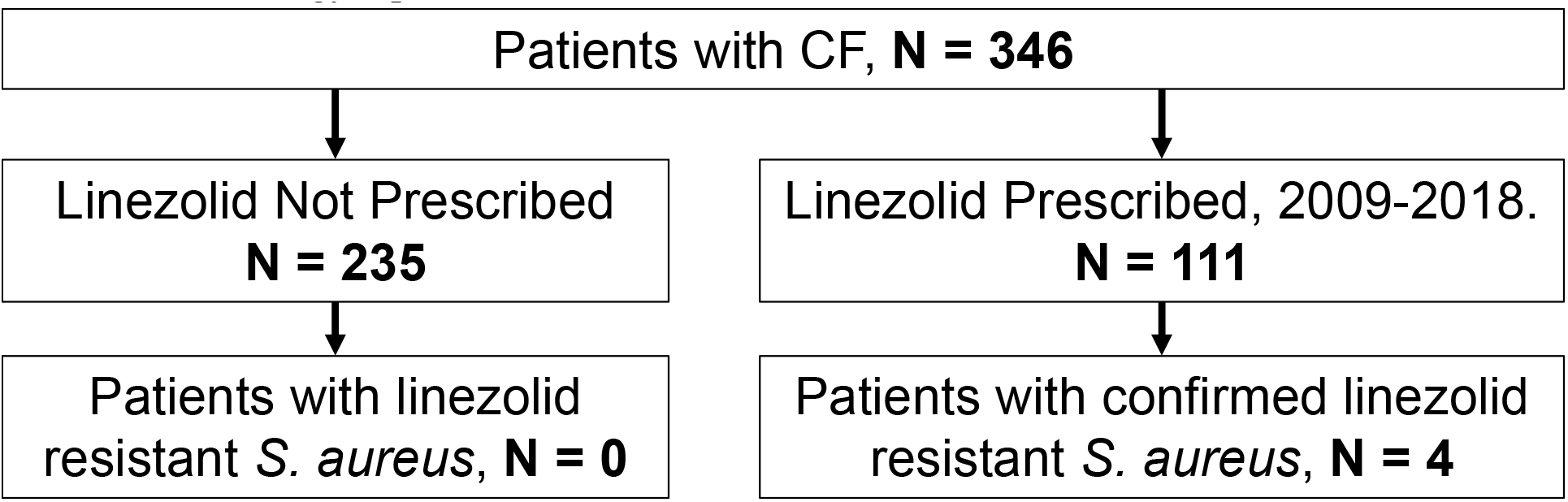
Eligibility Criteria. N = 360 patients with CF have records between 2008 and 2018. Of these, N = 346 had complete records including both electronic prescriptions and clinical microbiology reports.

**Supplemental Figure 3.**
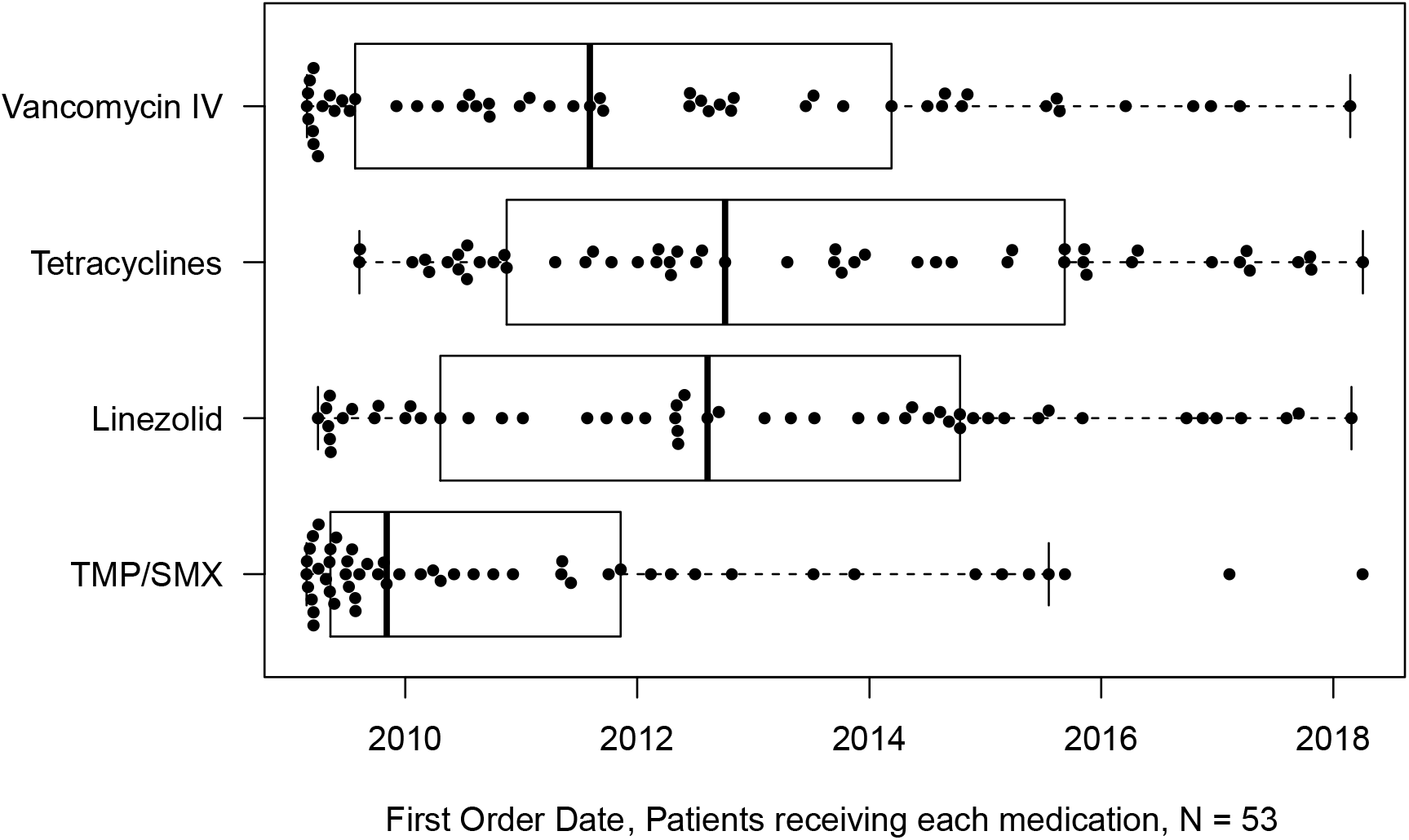
Sequence of administration of antibiotics active against MRSA. Linezolid and tetracyclines were used after prior attempts at treatment with trimethoprim-sulfamethoxazole (TMP/SMX) and Vancomycin IV. There were significant differences in the order of treatment, Friedman’s rank sum test *P* < 0.001. Linezolid followed vancomycin IV (Wilcoxon signed rank test *P* = 0.01) and TMP/SMX (*P* < 0.001).

**Supplemental Data 4.**
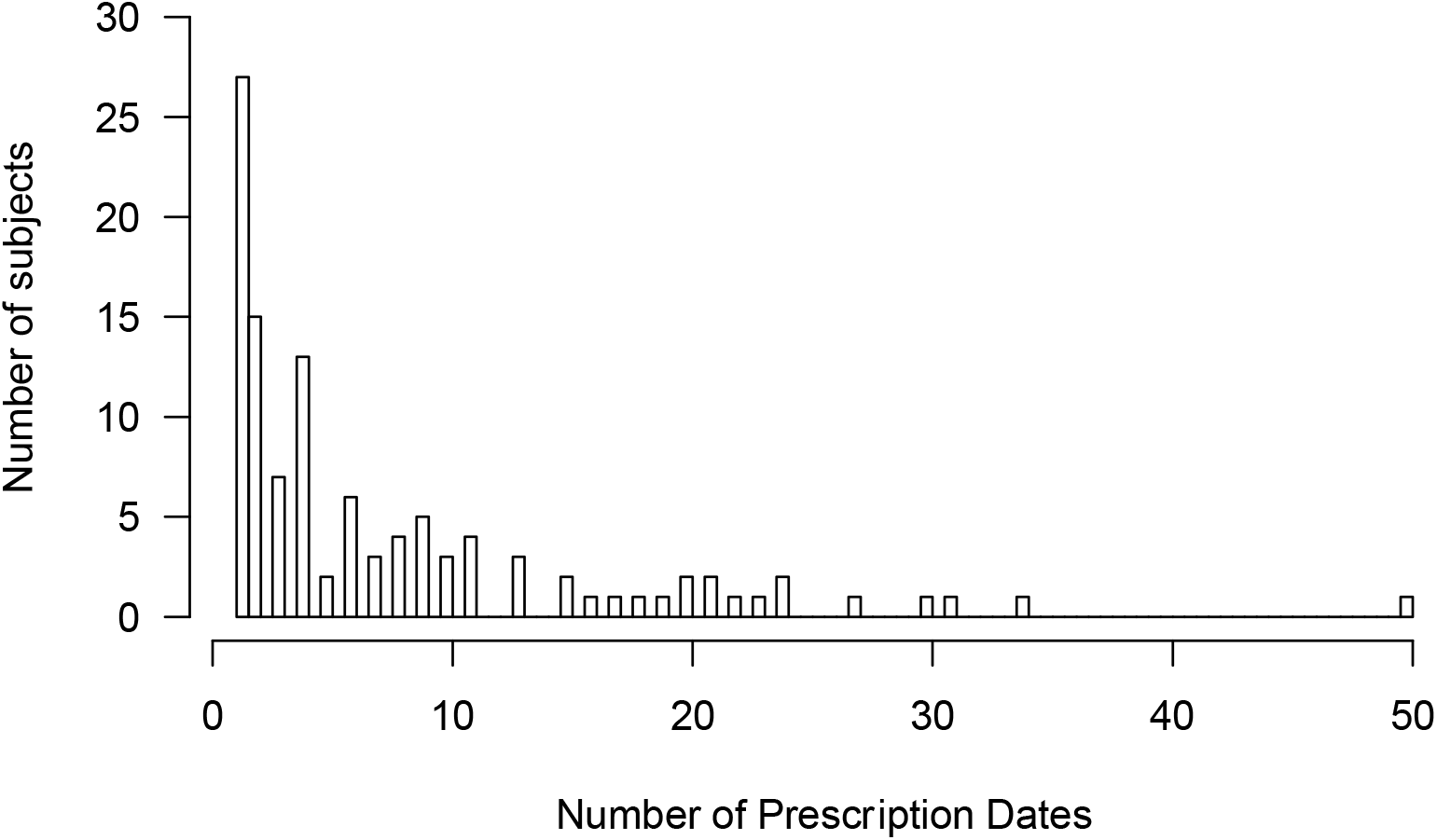
Number of unique dates with a linezolid order per patient.

**Supplemental Figure 5.**
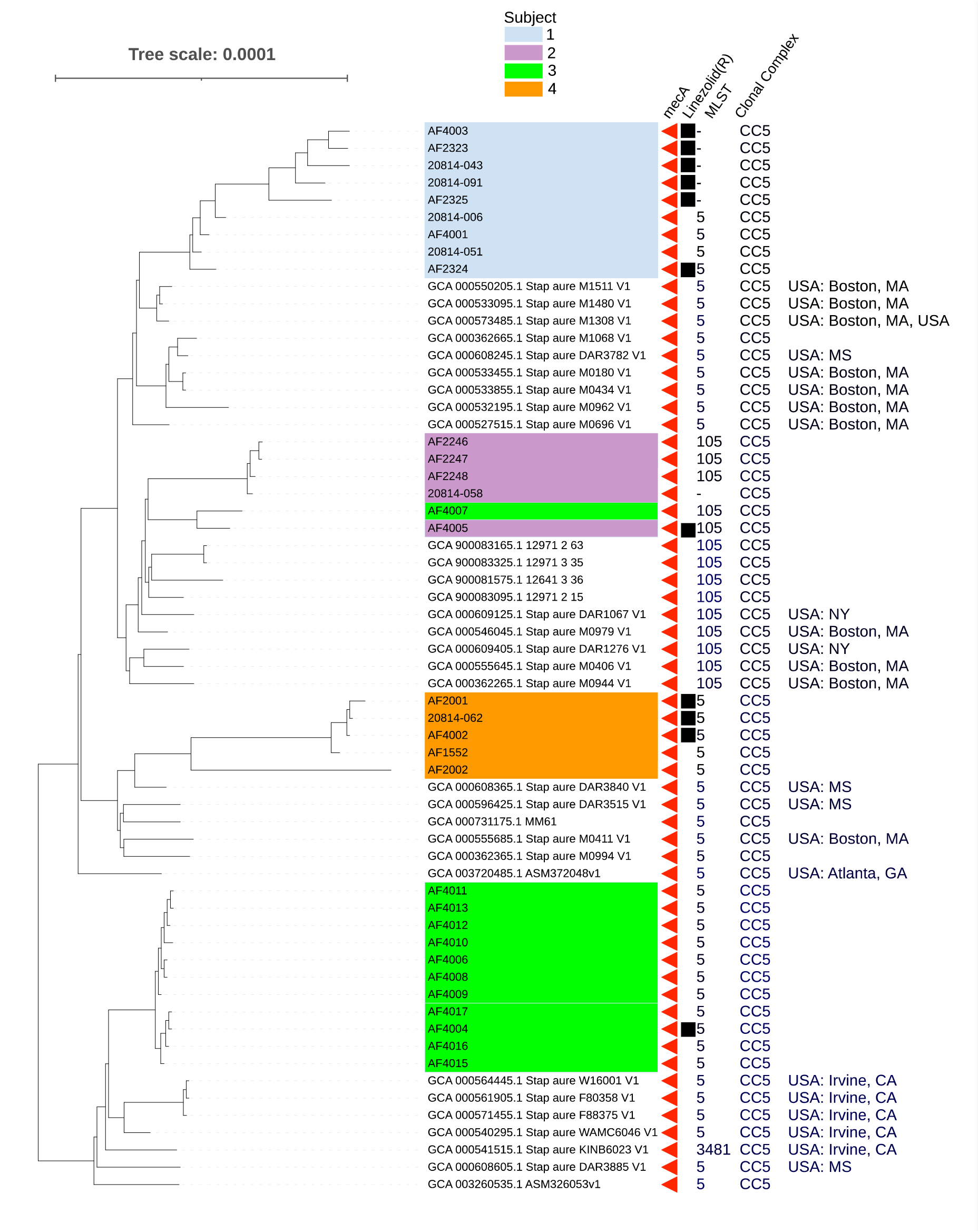
Phylogenetic tree based as in Figure 3, except branch lengths are drawn proportionate to nucleotide substitutions per site. For simplicity, we have omitted AF4014, an ST582 isolate. Five linezolid resistant isolates from Subject 1 (AF2325, HP20814-091, HP20814-043, AF2323, and AF4003) had complete loss of *mutS* and *mutL* associated with hypermutation, as shown by increased branch length. Similarly, AF2002 from Subject 4 had an increased number of substitutions; this isolate had a p.P340fs mutation from the loss of an adenine within a 9-adenine stretch.

**Supplemental Data 6.**
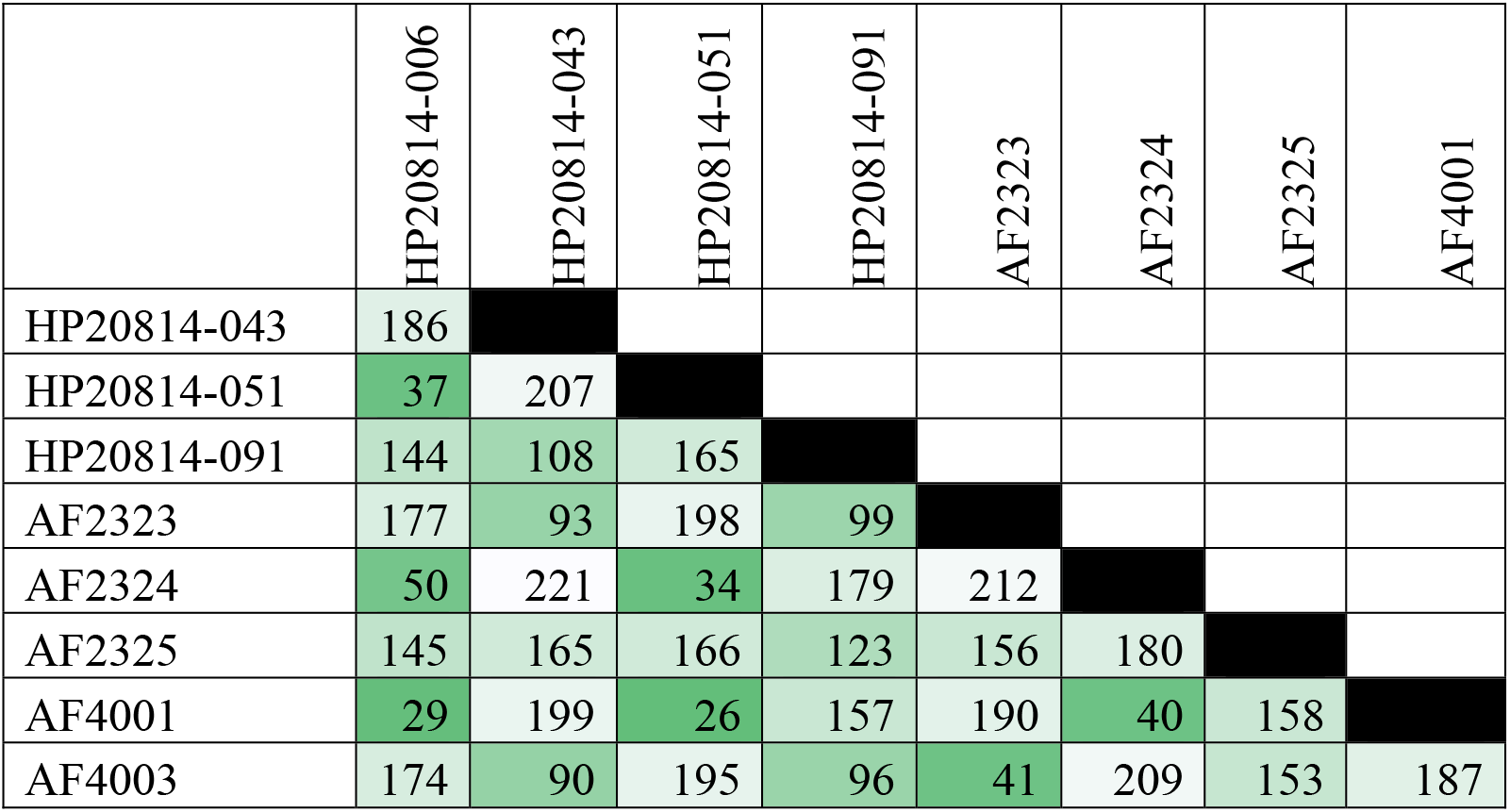
SNP distance between isolates from Subject 1. Several distances within this strain exceed the arbitrary limit of 60 SNPs. Black shading indicates identity. Darker green shading indicates greater proximity between isolates.

**Supplemental Data 7.**
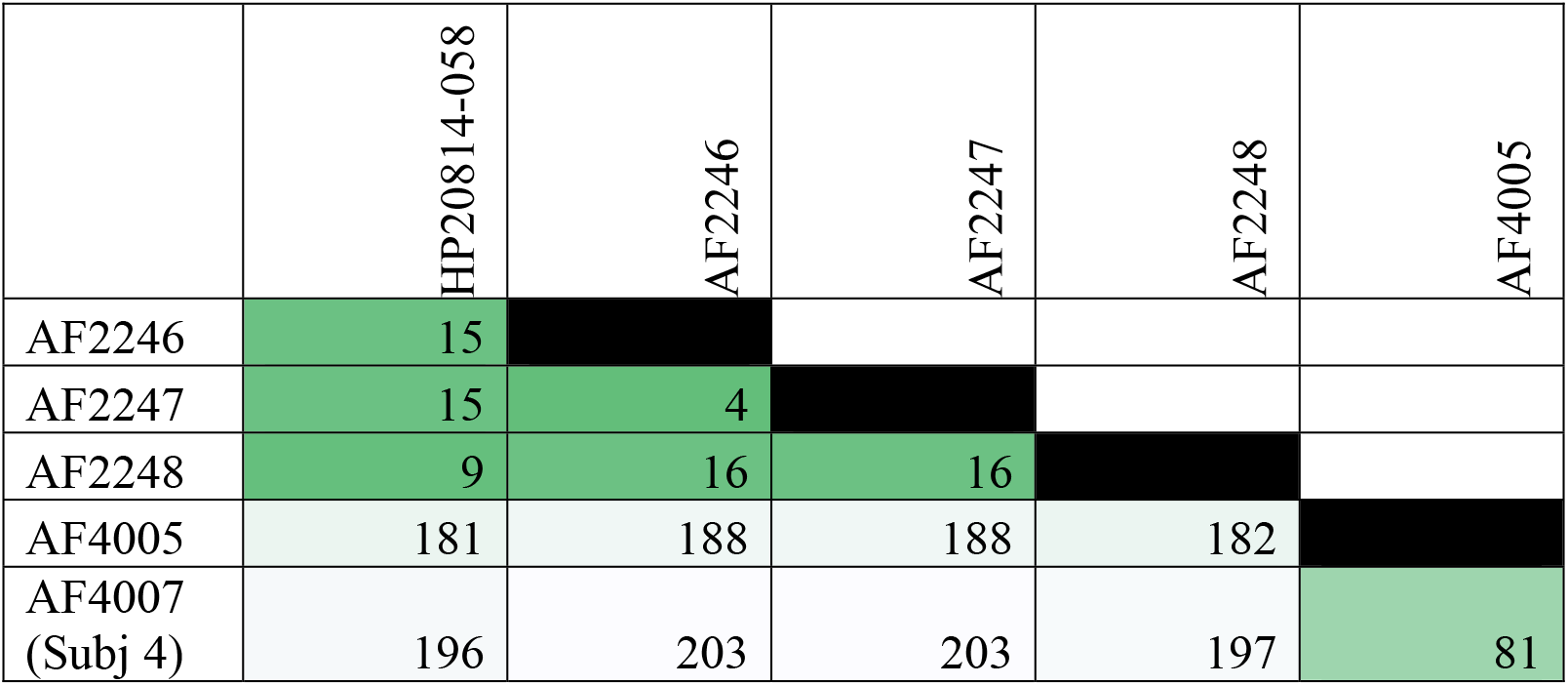
SNP distance between isolates from Subject 2 and shared strain AF4007 from Subject 4. Black shading indicates identity. Darker green shading indicates greater proximity between isolates. Although AF4005 and AF4007 are > 60 SNPs apart from other isolates, we cannot disprove that they belong to the same strain by Clade Breaker analysis.

**Supplemental Data 8.**
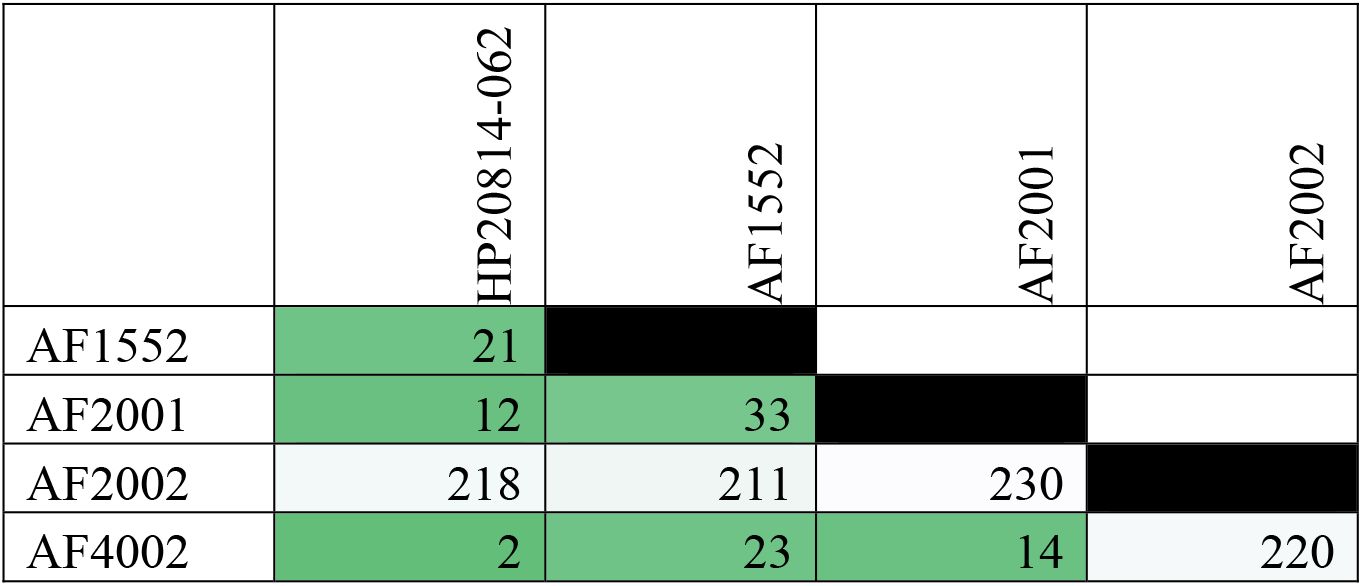
SNP distance between isolates from Subject 3. Black shading indicates identity. Darker green shading indicates greater proximity between isolates. AF2002 is the most distant, but we cannot disprove that it belongs to the same strain as the other isolates by Clade Breaker analysis.

**Supplemental Data 9.**
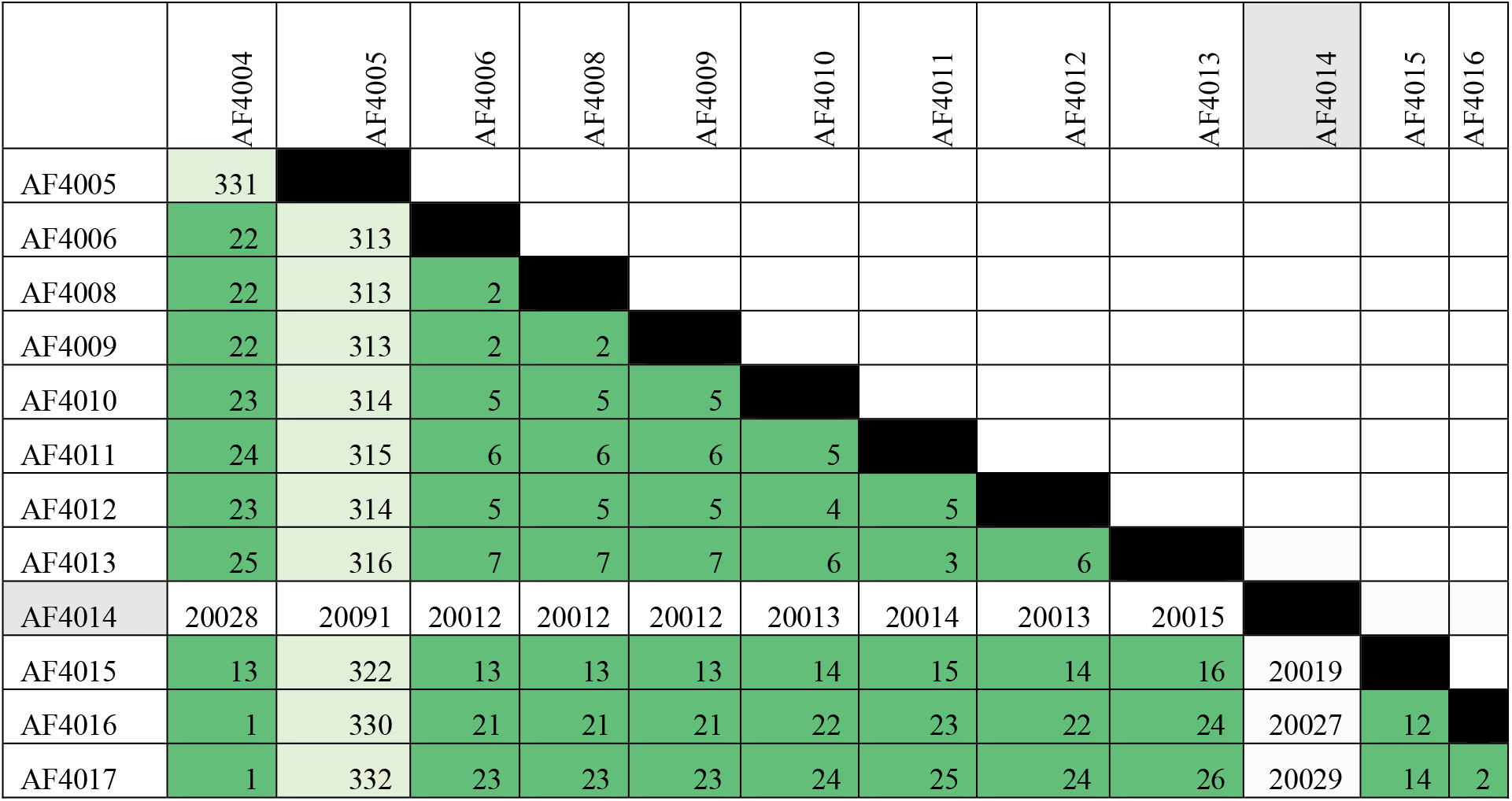
SNP distance between isolates from Subject 4. AF4014 is ST582. AF4005 belongs to a distinct strain of ST105, which has strong similarity to isolates from Subject 2. Black shading indicates identity. Darker green shading indicates greater proximity between isolates.

**Supplemental Data 10.**
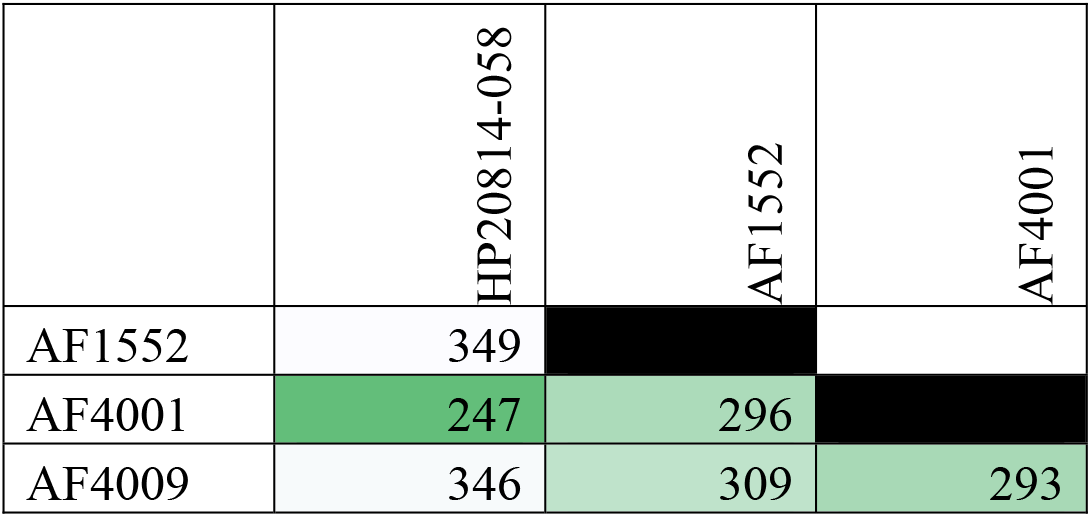
SNP distance between representative isolates from each subject.

**Supplemental Data 11.**
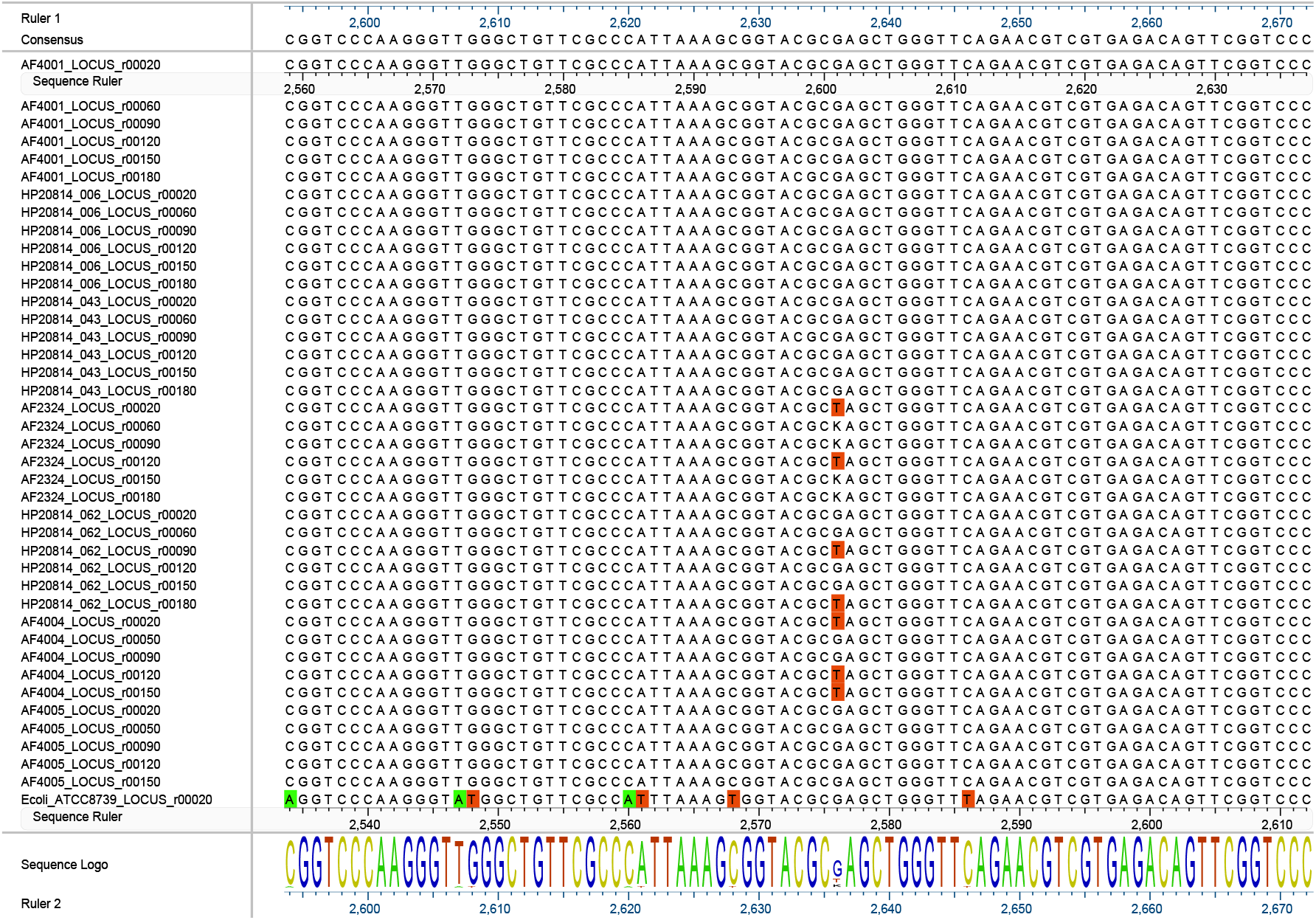
Repeated appearance of a known linezolid resistance mutation (23S rRNA G2576T) in unrelated isolates. Because of ambiguity in the numbering system for 23S rRNA genes (21), we used reference *Escherichia coli* strain ATCC 8739 (GenBank accession NC_010468) to indicate position 2576. AF2324 (Subject 1) contains 6 copies of 23S rRNA; 2 alleles were T, and the remaining reads were ambiguous (K indicates either G or T). AF4004 (Subject 3) contains 5 copies of 23S rRNA; three alleles are G2576T. HP20814-062 (Subject 4) contains 6 copies of 23S rRNA, 2 are G2576T.

**Supplemental Data 12.**
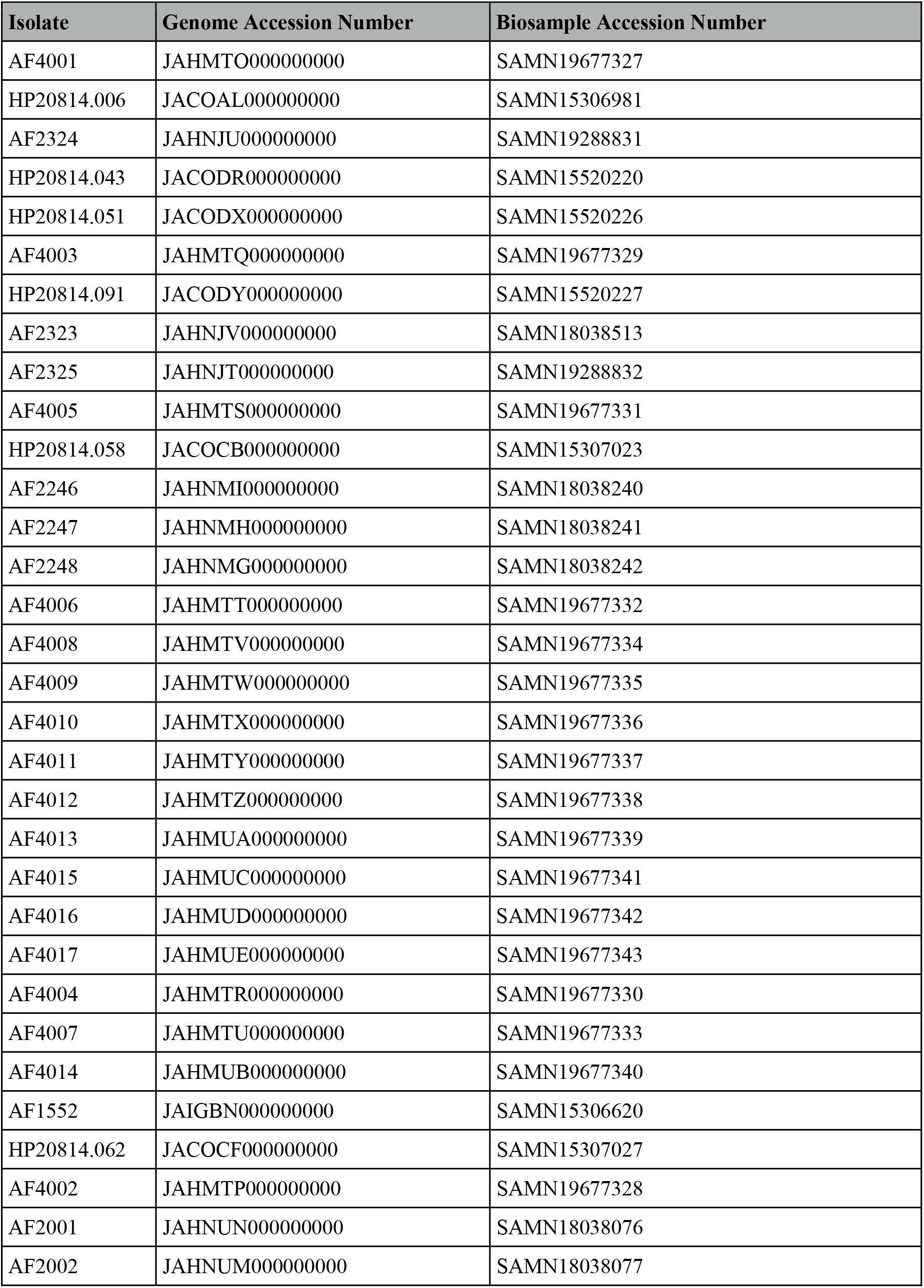
GenBank Accession numbers for isolates analyzed in this study.

